# Echo State Network models for nonlinear Granger causality

**DOI:** 10.1101/651679

**Authors:** Andrea Duggento, Maria Guerrisi, Nicola Toschi

**Author notes:** Corresponding author: A. Duggento.

## Abstract

While Granger Causality (GC) has been often employed in network neuroscience, most GC applications are based on linear multivariate autoregressive (MVAR) models. However, real-life systems like biological networks exhibit notable non-linear behavior, hence undermining the validity of MVAR-based GC (MVAR-GC). Current nonlinear GC estimators only cater for additive nonlinearities or, alternatively, are based on recurrent neural networks (RNN) or Long short-term memory (LSTM) networks, which present considerable training difficulties and tailoring needs. We define a novel approach to estimating nonlinear, directed within-network interactions through a RNN class termed echo-state networks (ESN), where training is replaced by random initialization of an internal basis based on orthonormal matrices. We reformulate the GC framework in terms of ESN-based models, our ESN-based Granger Causality (ES-GC) estimator in a network of noisy Duffing oscillators, showing a net advantage of ES-GC in detecting nonlinear, causal links. We then explore the structure of ES-GC networks in the human brain employing functional MRI data from 1003 healthy subjects drawn from the human connectome project, demonstrating the existence of previously unknown directed within-brain interactions. ES-GC performs better than commonly used and recently developed GC approaches, making it a valuable tool for the analysis of e.g. multivariate biological networks.

## I. INTRODUCTION

Multivariate Granger causality [1], [2] estimates how much the forecast of a timeseries can be improved by including information from the past of another timeseries, while accounting for additional, mutually interacting signals. It is defined in terms of conditional dependencies in the time or frequency domains [3], and can be considered an estimator for directed information flow between pairs of nodes (possibly) belonging to complex networks [4]. Granger Causality(GC)-based approaches, including the nonlinear Kernel approach[5] and the recent State Space (SS) (SS-GC) reformulation [6], have been employed in a vast number of problems which can be assimilated to network science, and the majority of CG applications are based on linear multivariate autoregressive (MVAR) models [2]. However, it is well known that real-life systems in general (and biological networks in particular) exhibit notable nonlinear behavior, hence undermining the validity of MVAR-based approaches in estimating GC (MVAR-GC) [7]. A typical case study is the analysis of brain networks from functional MRI (fMRI) signals, which result from convolving neural activity with a locally hemodynamic response function (HRF) [8], [9]. Here, a linear MVAR approach is not suitable for reconstructing neither the nonlinear components of neural coupling, nor the multiple nonlinearities and time-scales which concur to generating the signals. Instead, neural network (NN) models more flexibly account for multiscale nonlinear dynamics and interactions [10]. For example, multi-layer perceptions [11] or neural networks with non-uniform embeddings [12] have been used to introduce nonlinear estimation capabilities which also include “extended” GC [13] and wavelet-based approaches [14]. Also, recent preliminary work has employed deep learning to estimate bivariate GC interactions [15], convolutional neural networks to reconstruct temporal causal graphs [16] or Recurrent NN (RNN) with a sparsity-inducing penalty term to improve parameter interpretability [17], [18]. While RNNs provide flexibility and a generally vast modelling capability, RNN training can prove complex and their employment in real-world data, where data paucity is often an issue, may prove impractical and/or unstable. In this respect, a subclass of RNN, termed long-short term memory (LSTM) models, have been designed to explicitly include a “forgetting element” [19] which facilitates training (see [20] and references therein for a general discussion of LSTM in various learning tasks), and one paper also employed LSTM models in brain connectivity estimation [21]. Still, successful design and training of both RNN and LSTM models requires memory-bandwidth-bound computation, involves in-depth tailoring to a specific application, and the final architecture is often defined through trial and error procedures.

In this paper, we introduce a novel approach to estimating nonlinear, directed within-network interactions while retaining ease of training and a good degree of generality. Our framework is based on a specific class of RNN termed echo-state networks (ESN) [22]. The peculiarity of ESN is that, contrary to the general RNN model, ESN weights are not trained but rather randomly initialized, after which a linear mixing matrix is employed to map internal states to predicted outputs. The main hypothesis is that a fixed but randomly connected RNN can provide output with a state space rich enough to provide flexible fitting capabilities while eliminating the training issues common in RNNs. In addition, we modify and optimize the current ESN formulation to simultaneously model nonlinear, multivariate signal coupling while decoupling internal model representations into separate orthonormal weight matrices. We then reformulate the classical GC framework in terms of ESN-based models for multivariate signals generated by arbitrarily complex networks, and characterize the ability of our ESN-based Granger Causality (ES-GC) estimator to capture nonlinear causal relations by simulating multivariate coupling in a network of interacting, noisy Duffing oscillators. Synthetic validation shows a net advantage of ES-GC over other estimators in detecting nonlinear, causal links. As proof-of-concept, we then explore the structure of EC-GC networks in the human brain employed functional MRI data from 1003 healthy subjects scanned at rest at 3T withing the human connectome project (HCP), demonstrating the existence of previously unknown directed within-brain interactions.

## II. METHODS

### A. Granger causality

GC was introduced [23], [24] under the assumptions that a) the cause happens prior to its effect, and b) the cause contains unique information about the future of the effect. Under these assumptions, given a time-evolving system **u**(*t*) with *L* components {*u*_1_, *u*_2_, … *u*_*L*_}, component *u*_*j*_ is said to be causal on component *u*_*i*_ (*u*_*j*_ → *u*_*i*_) if:

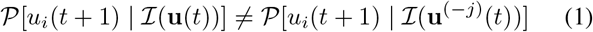

where 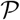 is a probability density function, while 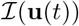 and 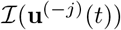 denote (with loose notation) all information provided by **u** up to time *t* including or excluding component *j*, respectively.

A common simplification is that 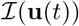 can be represented by a MVAR process defined over **u**. The inequality (1) is then replaced by a test of equality between the estimated variances of the two distributions, i.e. Var(*ε*′) ≠ Var(*ε*), where *ε* and *ε*′ are the prediction errors derived from the so called restricted model (RM) and an unrestricted model (UM), respectively:

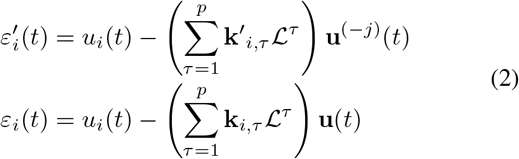

where 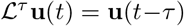 is the lag operator, the autoregressive order *p* is a suitably chosen parameter, and **k**′_*i,τ*_ and **k**_*i,τ*_ are to be estimated from data. Further, it is common practice to use the logarithm of the ratio of average squared residuals 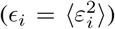 as a measure of MVAR-GC strength as follows: 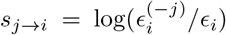 [1]. This measure also has a natural interpretation as the rate of “information transfer” between *i* and *j* and has been shown to be equal to the transfer entropy between *i* and *j* in the case of Gaussian variables [25]–[27]. Similarly, K-GC[5] is a nonlinear reformulation of GC based on searching for linear relations on a Hilbert space into which data has been embedded [28]. Also, more recently [6] the MVAR approach has been refined through a latent state-space (SS) model, where the inference of SS-GC is done over observables which are a linear mixture **v**(*t*) of the state variables **u**(*t*) with added white Gaussian noise: **v**(*t*) = *A***u**(*t*)+*ξ*(*t*) (where *A* is a mixing matrix). SS-GC has been extended in [29] to define multiscale causality, and has been shown to augment performance when classical MVAR methods fail [30].

### B. Echo State Network based causality

ESNs were introduced as a specific type of RNN which can be associated with an architecture and a supervised learning principle. In an ESN, a random, large and fixed RNN is fed with input signals eliciting a nonlinear response in each neuron within the network’s “reservoir”. The output is then derived as a linear combination of these nonlinear responses. In this way, the information 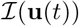 can be encoded through an *M* - dimensional array (the “reservoir”) of dynamical states **x**(*t*) which evolve in time as a function of the input **u** and of previous states **x**(*t* − 1). The reservoir is often [31] modeled with an exponentially decaying memory and an innovation terms 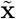 whose relative contribution is linearly weighted by the so called leak-rate *α*:

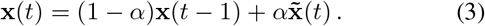

Additionally, the innovation term is a nonlinear function of the contribution of two other terms: a linear combination of the input states **W**^in^**u**(*t*), and a linear combination of the previous reservoir states **Wx**(*t* − 1), yielding 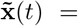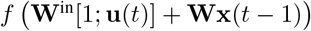. **W** ∈ ℝ^*M×M*^ is a mixing matrix between reservoir states, and **W**^in^ ∈ ℝ^*M×L*^ is a mixing matrix between input states. Typically, both **W** and **W**^in^ are constant and randomly initialized, and **W** is usually a sparse matrix whose initialization is controlled by its largest eigenvalue *ρ* (the so-called spectral radius) and its density. Also, a typical choice for the function *f*: ℝ^*M*^ → ℝ^*M*^ is an element-wise sigmoid function (e.g. hyperbolic tangent) which is symmetrical around the origin, approximates identity for “small” inputs, and is asymptotically bounded. The choices of *α*, **W**^in^ and **W** are crucial for forecasting accuracy [22].

### C. Redefining Causality trough ESNs

For a suitable choice of reservoir size *M*, leak parameter *α*, spectral radius *ρ*, matrix **W** and matrix **W**^in^, we can define an “extended” state **z**(*t*) = [1; **x**(*t*), **u**(*t*)] which, arguably, contains (most of) 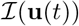. Then, under a linear approximation, we can assume that the expected value of 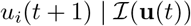 can be written as a linear combination of an “optimal” matrix **W**^out^ (to be estimated numerically) and **z**(*t*):

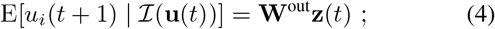

given a realization of the system, **W**^out^ ∈ ℝ^1×(1+*M*+*L*)^ is found by minimizing the sum of the squared residuals generated when using the next time-point of the *i*-th component as the ‘influenced’ variable. Then, the RM and UM in equation (2) can be reformulated as:

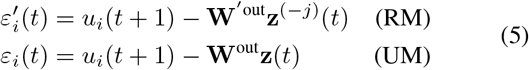

and, just like in the classical definition of GC, 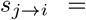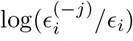 is the estimate of ES-GC strength.

In this paper, under the assumption that each component of **u** interacts weakly (as compared to its own dynamics) with other components, we introduce the choice of **W** as a block diagonal matrix **W** = diag(**W**_1_, **W**_2_, …, **W**_*L*_). Equivalently, we assume that, for each component *i*, the expected value of 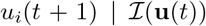 is linearly separable in terms of all echo states including its own. Figure 1 shows a pictorial representation of this model, which can also be thought of as a larger ESN composed of several separable ESNs. As a further improvement, in this paper we introduce the use of orthonormal matrices (as opposed to sparse, randomly initialized matrices) as the block diagonal matrices **W**_*i*_, obtained through random initialization followed by orthonormalization. This is heuristically motivated by the idea of providing the network with a “maximally orthogonal” basis for signal representation. Experimentally, we found that this choice i) consistently yields superior forecasting performance in terms of residual sum of squares of univariate models, and ii) renders performance largely insensitive to the choice of parameters (*ρ,* **W**^in^) within a wide range of values (data not shown).

**Figure 1.**
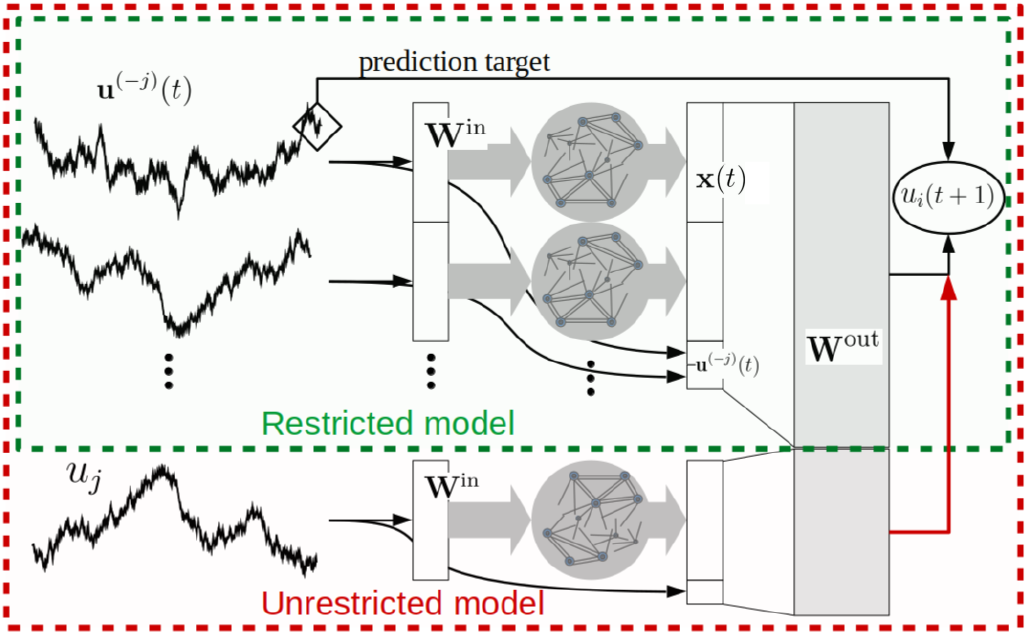
Schematic representation of ES-GC.

### D. Synthetic validation of ES-GC and comparison to other estimators

#### 1) Network generation

In order to compare the performances of ES-GC, SS-GC, K-GC and MVAR-GC in detecting true causal connections within complex directed networks, we generate data from a family of 10-node ground-truth random networks derived by the Erdös-Rényi model [32], [33]. This entails randomly sampling from a uniform graph distribution, i.e. a graph is constructed by connecting nodes randomly or, equivalently, each edge is included in the graph with constant probability independent from every other edge. Specifically, starting with *L* disconnected nodes, “edges” (i.e. connections) between two not already connected nodes are successively and randomly assigned up to the required density. Bidirectional connections as well as loops are explicitly allowed. The total number of edges *n*_e_ depends on the network density *d*_n_ which, for a network with *L* nodes, is defined as *n*_e_/(*L*(*L* − 1)). Here, we generated graph families at 9 different densities, where values are chosen so that the corresponding densities are approximately equidistant on a logarithmic scale between 0.01 and 1: *d*_n_ = {0.022, 0.044, 0.067, 0.1, 0.154, 0.249, 0.387, 0.584, 0.822}.

For each value of *d*_n_ we generate 30 different networks to account for fluctuations with respect to network topology. Each network is described by a binary, zero-diagonal, asymmetric adjacency matrix **A**, whose elements *A*_*ij*_ represent the direct influence of node *j* on note *i*. Examples of the generated networks at different densities are shown in Fig. 2.

**Figure 2.**
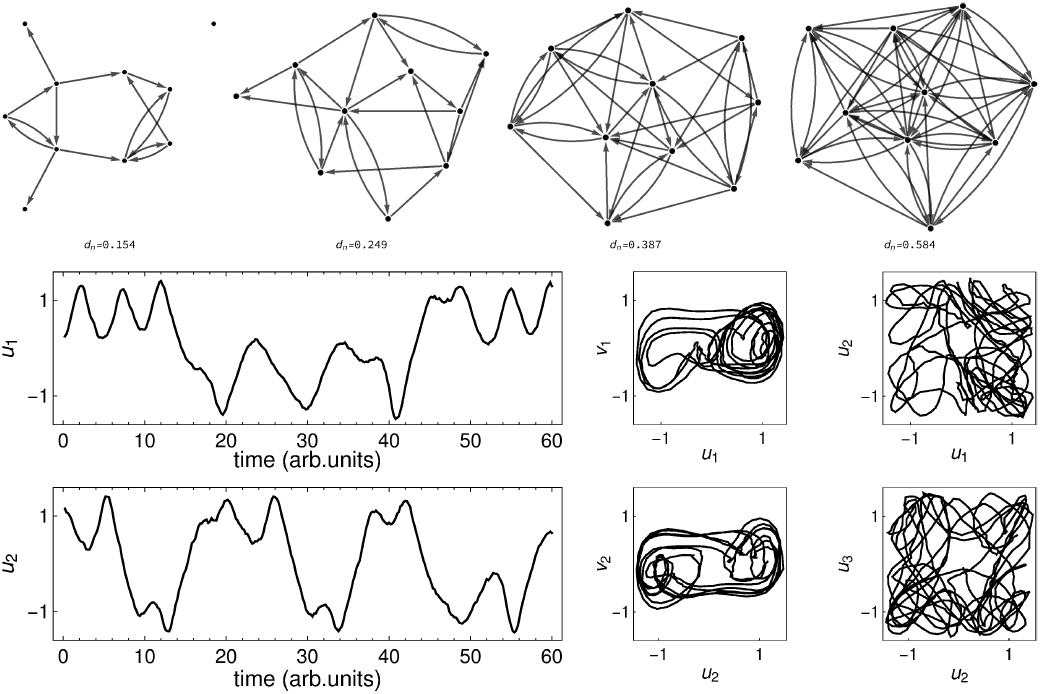
Top: example networks used to generate synthetic data at densities *d*_n_ = 0.1544, 0.2485, 0.387, 0.5837. Bottom: signals from a network of forced, weakly coupled Duffing oscillators (eq.(6)). Parameters were chosen so that: i) each oscillator would exhibit chaotic behaviour even without coupling; ii) none of any two oscillators would be synchronized regardless of network density or coupling strength. In this paper, *γ* = 0.5, *β* = −1, *α* = 1, *δ* = 0.3; *ω*_*i*_ is randomly chosen from the interval [1.19; 1.21] and *ϕ*_*i*_ is randomly chosen from the interval [0; 2*π*]; Σ = *σ*^T^*σ* where *σ* = **m**^T^(0.05 *I*)**m** and **m** is a matrix whose elements are randomly sampled from a uniform distribution in the [−1;1] interval.

#### 2) Node-wise Duffing oscillators

For each ground-truth network, a set of forced, noisy, weakly coupled, Duffing oscillators **u** = {*u*_1_, …, *u*_*L*_} are generated and assigned to network nodes as follows:

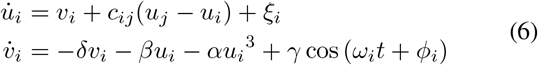

where *ξ*_*i*_(*t*) is a spatially correlated white noise process: ⟨*ξ*_*i*_(*t*), *ξ*_*j*_ (*τ*)⟩ = *δ*(*t* − *τ*)Σ_*ij*_. The coupling coefficient *c*_*ij*_ is defined by a global coupling strength *w* and a ground-truth matrix **A** that defines the topology of the network. Specifically, for the *i*-th node, if *A*_*ij*_ = 0 then *c*_*ij*_ = 0; otherwise *c*_*ij*_ is equal to *w* normalized by the number of incoming connections *c*_*ij*_ = *w*/Σ_*i*_*A*_*ij*_. Here, nine values (approximately equidistant on a logarithmic scale) for *w* were employed (*w* = 0.02 − 0.5). Each networks was evolved for a total of 10000 timepoints (Δ*t* = 0.5, undersampled from a signal generated with a stochastic second-order Runge-Kutta numerical integration scheme with integration step Δ*t*′ = 0.1). Example synthetic signals are shown in Fig. 2 along with details about parameter choices.

#### 3) Causality estimation in ground truth networks

For each set of synthetics signals (30 networks × 9 density values × 9 coupling strengths = 2430 networks with 10 nodes each), the four estimation methods (ES-GC, SS-GC, K-GC and MVAR-GC) were employed for ground-truth network reconstruction. For each {*w*, *d*_n_} pair, the performance of each estimator was quantified through a receiver operating characteristic (ROC) curve built by varying the threshold in causality strength used for edge acceptance across all network edges. Performance metrics derived at each density and coupling strength were averaged across the 30 networks. ^1^.

For both SS-GC and MVAR-GC, the optimal autoregressive order *p* was chosen according to the Akaike information criterion (AIC) within the range 1-25 [34] (*p* = 20 for both estimators). The optimal *p* was not significantly sensitive to network density (data not shown). ES-GC hyperparameters were chosen by performing a 3-1 train-test split on univariate data generated from one network node and minimizing prediction error on the test set as a function of reservoir size *M* (interval: 1-500), leak-rate *α* (interval: 0.1-0.9) and spectral radius *ρ* (interval: 0.01-1) [35]. This procedure was repeated for varying data length (interval 2048-65536) This resulted in *M* = 2500 (corresponding to a reservoir of 250 neural units for each of the 10 nodes), *α* = 0.3 and *ρ*=0.9.

The area under the ROC curve (AUC) was employed as a performance metric. Since SS-GC is an explicitly multi-scale method [29], SS-GC estimations were repeated at 18 different scales (1-18). In this paper, all AUC values presented for SS-GC are the highest value achieved amongst all scales. A similar procedure was followed for K-GC, where estimations were repeated while concurrently varying model order (interval: 1-7) and polynomial kernel order (interval: 1-7). All AUC values presented are the highest value achieved within this parameter space. Additionally, we evaluated detection performance of all causality estimators in terms of the positive predictive value (PPV) of the top 10% strongest connections.

### E. Estimation of the human between-network connectome from fMRI data

As an example application to biological data, we use in-vivo fMRI data from 1003 subjects made available by the Human Connectome Project [36] as part of the S1200 PTN release. The subjects included underwent 4 sessions of 15-minute multi-band (repetition time (TR) = 0.72s) resting-state fMRI scans on a 3 Tesla scanner with isotropic spatial resolution of 2 mm, for a total of 4800 volumes per subjects. Preprocessing details can be found in [37]. After pre-processing, a group-principal component analysis [38] output was generated and fed into group-wise spatial independent component analysis (ICA) using FSL MELODIC tool [39] to obtain 15 distinct spatiotemporal components. Subject- and components- specific timeseries were then extracted, and a directed connectome was built for each subject through our ES-GC method. ES-GC hyperparameters were chosen as described above (using a train-test split of ICA-timeseries data), resulting in *M* = 60 (corresponding to a reservoir of 4 neurons for each of the 15 components) leak-rate *α* = 0.6, and spectral radius *ρ*=0.9. Interestingly, we obtained a smaller optimal reservoir and a larger optimal leak-rate as compared to the synthetic data case, possibly indicating less rich ‘dynamics’ in fMRI data as compared to networks of nonlinear duffing oscillators.

## III. RESULTS

### A. Synthetic validation results

Figure 3 shows the comparison between the ROC curves obtained when using ES-GC, SS-GC, K-GC and MVAR-GC for exemplary density and coupling parameters *d*_n_ = 0.154 and *w* = 0.1. ES-GC clearly outperforms SS-GC (even at its optimal scales), K-GC and MVAR-GC (which only delivers chance-level performance). Additionally, the ROC curves show how for ES-GC true positive rates/false positive rates are larger/smaller (respectively) than for other estimators at every discrimination threshold (i.e. operating point of the ROC curve).

**Figure 3.**
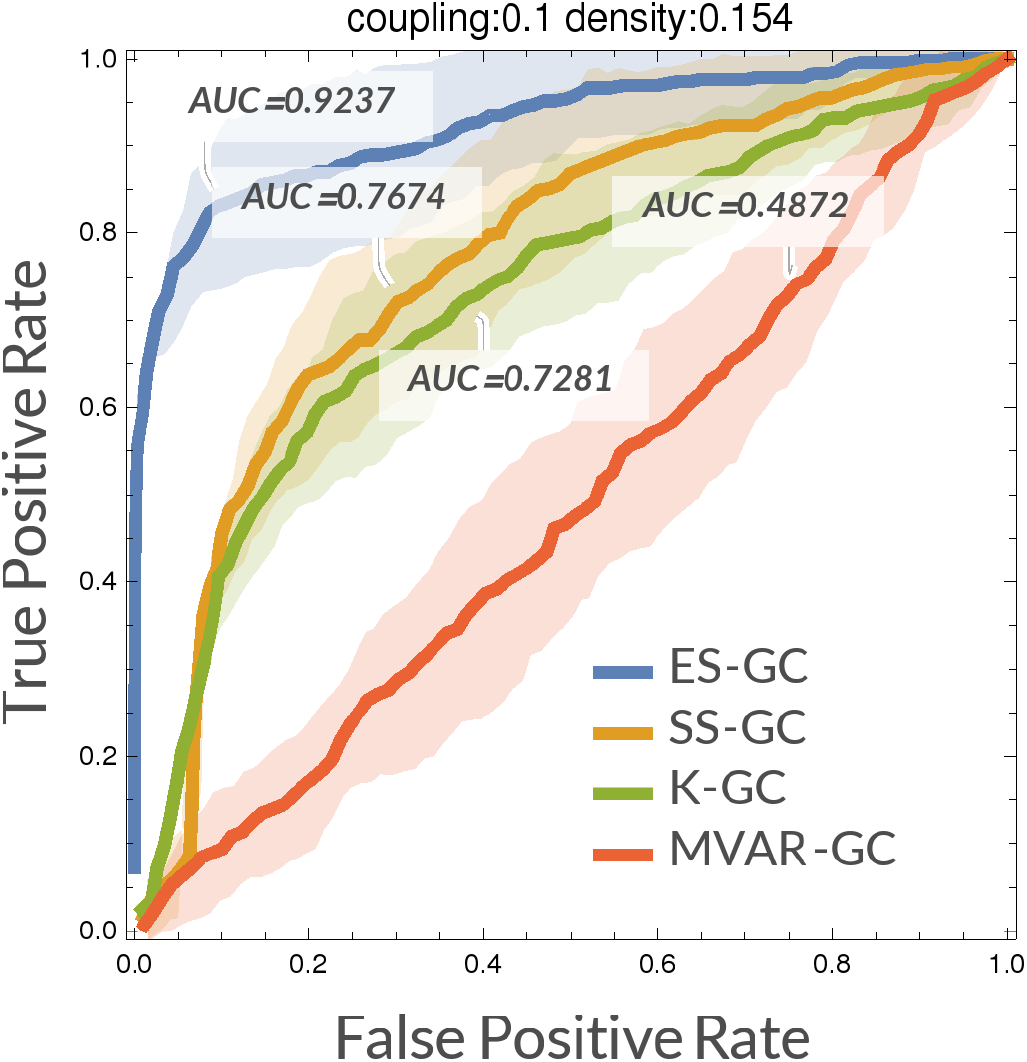
ROC curves relative to ES-GC, SS-GC, K-GC and MVAR-GC. ROC curves (built over the prediction of (10^2^ − 10) links for each of 30 networks and successively averaged) are shown (solid lines) along with a ± 1 standard deviation interval (shaded areas). ROC curves are show for parameters: density *d*_n_ = 0.154, coupling strength *w* = 0.1. All other parameters are kept constant (see Fig. 2).

Since performance of any causality estimator is expected to increase with coupling strength and to fluctuate with network density, we inspected AUC as a function of *w* with fixed *d*_n_ and vice versa (Figure 4). For all estimators, the AUC increases with coupling strength (a higher coupling corresponds to a larger multivariate transfer entropy [25] up to the onset of generalized synchronization (data not shown)). ES-GC performs notably better than other estimators at all network densities and all coupling strengths. Also, Figure 5 shows the comparison in PPV (for the top 10% strongest connections) between all frou estimation methods. For all estimators, PPV increases both with coupling strenght and with density. Again, ES-GC delivers notably higher PPV than other estimators at all network densities and all coupling strengths.

**Figure 4.**
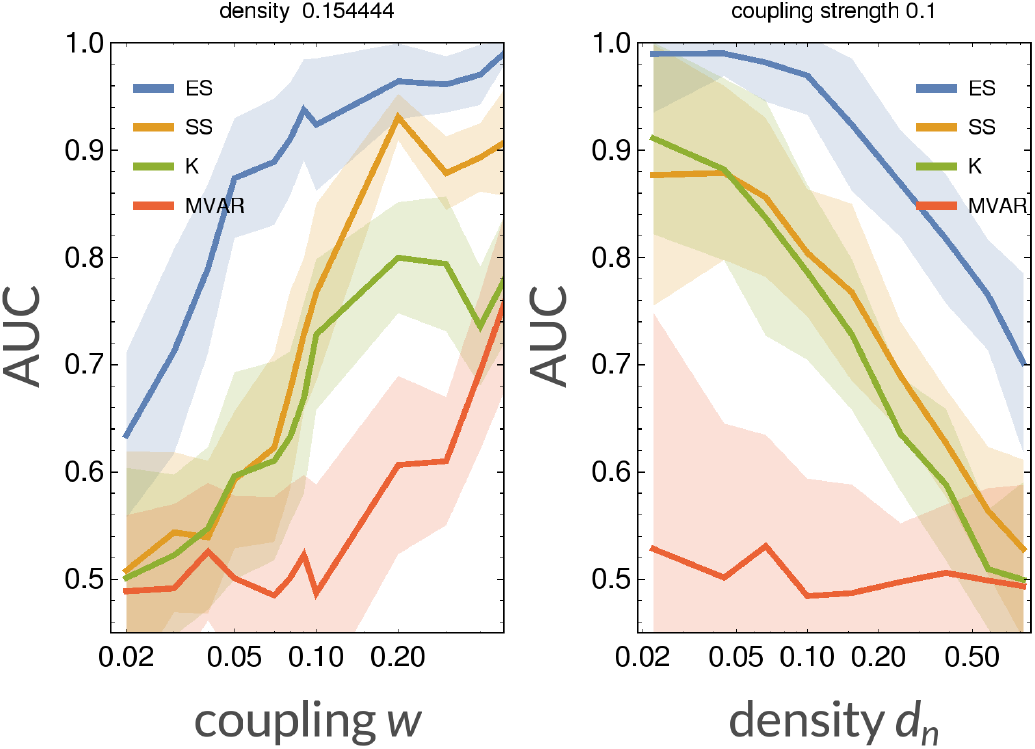
AUC comparison between ES-GC, SS-GC, K-GC and MVAR-GC with respect to coupling strength (left) and network density (right). For each method, AUCs were computed from ROC curves built over the prediction of (10^2^ − 10) links for each of 30 networks and successively averaged (solid lines). A ± 1 standard deviation interval is also shown as shaded areas.

**Figure 5.**
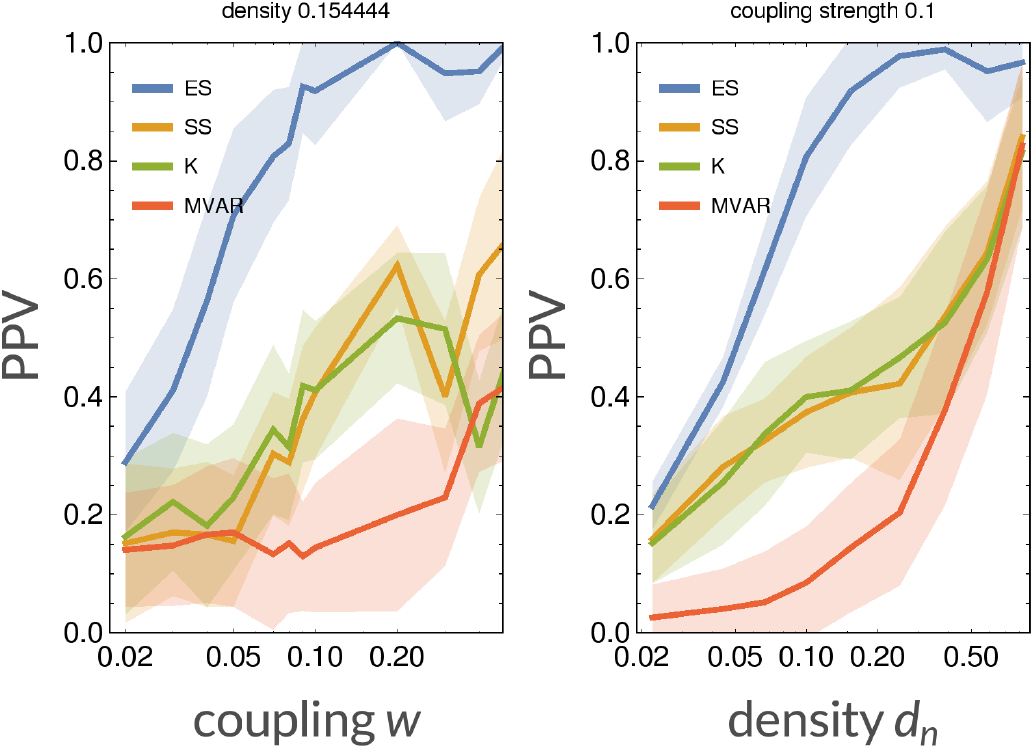
PPV of the top 10% connections between ES-GC, SS-GC, K-GC and MVAR-GC with respect to coupling strength (left) and network density (right). For each method, the PPVs built over the prediction of 30 different networks were averaged (solid lines); for each method shaded areas indicate the mean PPV ± 1 standard deviation.

#### In-vivo human connectome results

ES-GC estimation in the full HCP sample resulted in 4 × 1003 = 4012 asymmetric adjacency matrices. For the purpose of visualization (see below), we calculated the element-wise median matrix across subjects and scans, which was then thresholded at the 90th percentile. The resulting directed, within-component connectome derived from 1003 healthy subjects is shown in Fig. 6 (see Figure caption for the physiological significance of each of the 15 components). These results suggest a strong bidirectional interaction between the Default Mode Network and the Salience network, a direct modulation of the Striate Visual Network by the Visuo-Prefrontal Network (but not vice versa) and a direct modulation of the Hippocampal-Cerebellar Network by the Sensory/Motor-Limbic-Network, which was only recently defined [40].

**Figure 6.**
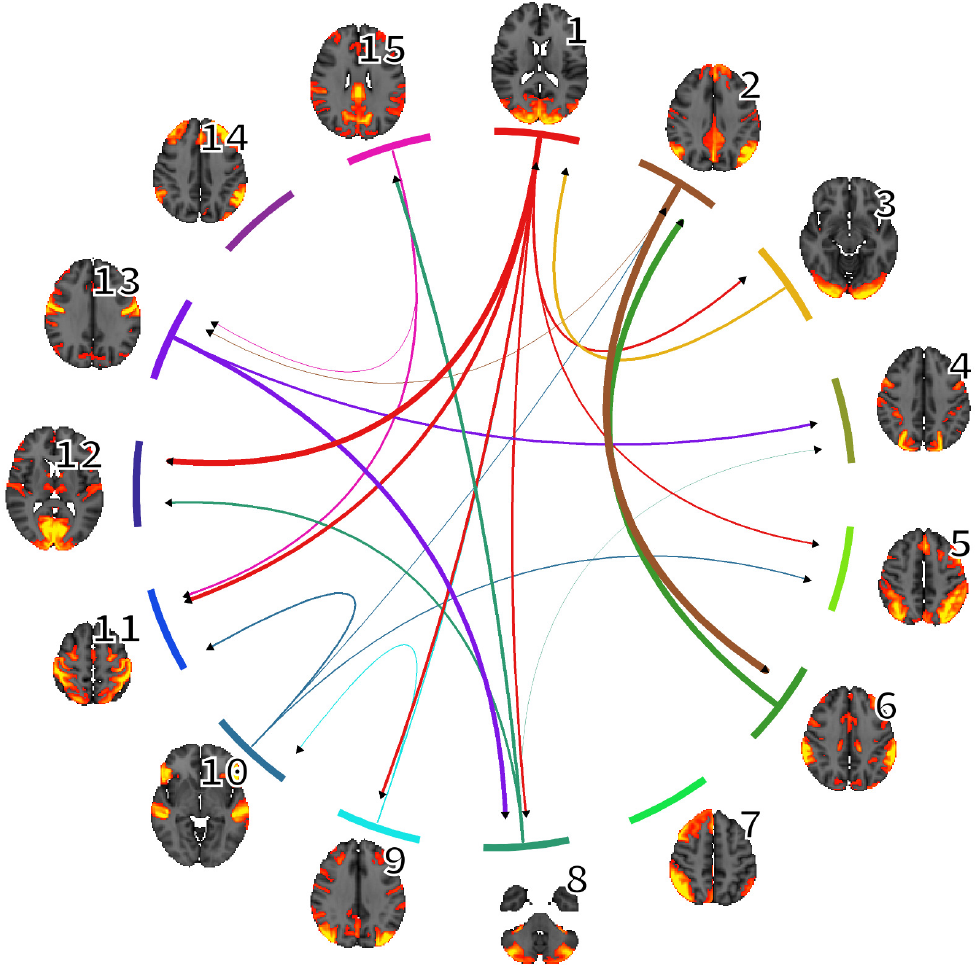
Graphical summary of in-vivo results. Directed, between-component brain connectivity network derived in 1003 HCP subjects (top 10% in median across subjects) is displayed. Color indicates the ‘influencing’ component. The significance of physiological networks can be summarized as follows [40]: 1: Visuo-Prefrontal Network, 2: Default Mode Network, 3: Extra-striate Visual Network, 4: Visuo-Parieto-Premotor Network, 5: Left Fronto-Parieto-Cerebellar Network, 6: “Salience” Network, 7: Right Fronto-Parieto-Cerebellar Network, 8: Hippocampal-Cerebellar Network, 9: Hippocampal-Posterior Cingulate Network, 10: Fronto-Temporal-Network, 11: Sensory-Motor Network, 12: Striate Visual Network, 13: Sensory/Motor-Limbic Network, 14: Fronto-Polar Network, 15: Cingulate Cortex Network.

## IV. DISCUSSION AND CONCLUSIONS

In this paper we transition away from classical causality quantifiers and define a novel approach to estimating nonlinear, directed network interactions through a specific class of RNNs (namely echo-state networks, or ESN) which do not suffer from training difficulties common in RNNs. We modify the current ESN formulation to represent nonlinear, multivariate signal coupling while decoupling internal model representations and using separate, heuristically motivated orthonormal bases for network weights. We then reformulate the classical GC framework in terms of ESN-based models for multivariate signals generated by complex networks. Our method is validated through extensive synthetic data simulation, where we find that ES-GC largely outperforms state-of-the art linear methods. Interestingly, in our model system, the coupling value *w* = 0.1 marks a transition across chance-level performance for classical MVAR-GC. This indicates that ESN-, K- and SS-based GC are capable of a non-random resolution of true links in a system where coupling is so weak that standard MVAR-GC analysis fails. Also, for all methods, performance mostly deteriorates with increasing network density. This is possibly due to the complex interplay between the dynamics of a high number of ‘causal’ nodes. Still, across the whole parameter space and model system parameters tested in this paper, ES-GC performs notably better than other methods. Importantly, these overall considerations also hold when looking at the PPV achieved when predicting the top 10% strongest connections, indicating that in possible applications (like e.g. the approach we adopted in this paper when deriving our proof-of-concept directed connectome) targeted at interpreting the strongest links ES-GC would deliver the best true-positive rate. While these results have been derived using forced, weakly coupled stochastic oscillators, the generality of the ES-GC framework allows to hypothesize that it could deliver superior performance in a larger class of dynamical situations like e.g. non-synchronized, weakly interacting, possibly chaotic and forced dynamical systems, in which causality detection is extremely challenging. Importantly, these circumstances are ubiquitous in biophysics and biomedical signals, where the detection of causal links in weakly interacting systems is often the stepping stone for physiological interpretation. In this context, our method was able to uncover direct functional links between sub-networks of the brain which have not been previously described. Relatedly, while the fMRI results in this paper are intended to demonstrate a possible application of our novel methods to real-world brain data, it is interesting to note the application of GC in neuroscience in general have been the object of constructive discussions [30], [41]–[44] which ultimately confirmed its applicability provided possible methodological pitfalls are avoided. For example, it is well known that fMRI signals are a surrogate of neuronal activity, and that the convolution with a locally varying HRF can confound causality results. Blind deconvolution methods have been proposed in this respect [45]. Still, the application of this type of pipeline is not yet widespread in fMRI studies using causality methods, and the importance and applicability of such methods in cases where fMRI time-series data is averaged over relatively large brain regions stemming from low-dimensional independent component analysis (line in this paper) remains to be investigated. Also, accurate synthetic simulations of neuronal spiking and neurovascular coupling have shown that the top percentiles in the median causal adjacency matrix computed across subjects can be interpreted with extremely good positive predictive value [9]. Relatedly, it has been shown that GC in general is applicable to fMRI data and that HRF convolution retains monotonicity between fMRI causality and neural causality at realistic fMRI temporal resolution and noise level [46], which corroborates the idea of reliably interpreting the top percentiles of causal connections found in a large number of subjects. Also, the importance of accounting for nonlinearity in GC estimates in fMRI (which is included in our model) as well as the problem of non-overlapping regions of interest across subjects (which is circumvented by employing group-ICA) and of employing non-equilibrium timeseries (which, however, does not apply to independent components derived from resting spate data like in this paper) have been previously underlined [47].

In summary, ES-GC performs significantly better than commonly used and recently developed GC detection tools, even in complex networks with nonlinear signals and weak coupling, making it a valid tool for the analysis of e.g. multivariate biological networks. Future work will address the incorporation of structural priors from e.g. diffusion MRI [48] as well as the reformulation through recent, more sophisticated NN architectures (e.g. combinations of combination of ESN and LSTM models [49]) which have been built to facilitate training.

## AUTHOR CONTRIBUTIONS STATEMENT

Conceptualization, AD and NT; Methodology, AD and NT; Formal analysis, AD; Funding acquisition, NT and MG; Investigation, AD; Supervision, NT; Writing – original draft, AD and NT; Writing – review and editing, AD, MG, NT. All authors critically reviewed the initial manuscript and approved the final manuscript as submitted.

## ADDITIONAL INFORMATION

The authors declare no competing interests, or other interests that might be perceived to influence the results and/or discussion reported in this paper.

## Footnotes

1 In-house developed code for ES-GC estimation is available on GitHub repository: https://github.com/andreaduggento/EchoState-GrangerCausality

